# Enhancing cortical electrophysiological signatures of sleep overrides ER stress responses in sleep-deprived mice

**DOI:** 10.64898/2026.10.16.759564

**Authors:** Jose Prius Mengual, Anna Hoerder-Suabedissen, Rong Mao, Lewis Taylor, Atreyi Charkrabarty, Sarah E Newey, Aarti Jagannath, Colin J. Akerman, Vladyslav V. Vyazovskiy

**Author notes:** <u>Corresponding authors</u>: Jose Prius Mengual, Department of Physiology, Anatomy and Genetics, Sleep and Circadian Neuroscience Institute (SCNi), Kavli Institute for Nanoscience Discovery Sherrington Rd, Oxford OX1 3QU, Vladyslav V. Vyazovskiy, Department of Physiology, Anatomy and Genetics, Sleep and Circadian Neuroscience Institute (SCNi), Kavli Institute for Nanoscience Discovery Sherrington Rd, Oxford OX1 3QU.

## Abstract

Sleep deprivation (SD) triggers characteristic changes in the cortical electroencephalogram (EEG) during waking and sleep, as well as upregulating molecular signatures of neuronal activity, synaptic plasticity, and endoplasmic reticulum (ER) stress. This includes the induction of the unfolded protein response (UPR), the primary marker and adaptive mechanism of ER stress. Yet how cortical neuronal activity during SD contributes to the induction of these molecular signatures remains unknown. Here we investigate whether manipulating cortical activity during SD with a physiological stimulation paradigm (flickering light stimulation, FLS) affects the induction of the UPR. As expected, SD alone upregulated activity-dependent and UPR-related genes in cortex and led to increased EEG slow-wave activity (SWA) during subsequent NREM sleep, an electrophysiological signature of sleep need. Delivering FLS during SD, however, prevented the upregulation of activity- and UPR-related genes while inducing an additional increase in SWA during subsequent sleep. Further analysis of neuronal firing revealed that FLS induced brief neuronal silent (OFF) periods in awake animals during stimulation, thereby shifting cortical activity towards a brain state resembling sleep. These findings suggest that physiological manipulation of cortical activity during waking is sufficient to dissociate molecular signatures of ER stress from electrophysiological markers of sleep pressure. We propose that global sleep-wake history and cortical network dynamics differentially regulate activity-dependent transcription and the accumulation of sleep pressure.

## Introduction

Sleep is essential for normal cognitive function, including attention, memory, and behavioural performance. Prolonged wakefulness impairs cognition and alters cortical activity in both humans and animals (Andrillon et al., 2021; Huber et al., 2004; Nir et al., 2017; Van Dongen, 2025; Vyazovskiy et al., 2011). However, the biological role of sleep in general and the specific physiological characteristics defining sleep largely remain elusive. One of the electrophysiological markers of sleep homeostasis is electroencephalographic (EEG) slow-wave activity (SWA; 0.5–4 Hz) during non-rapid eye movement (NREM) sleep, which increases as a function of prior wake duration and declines during subsequent sleep (Borbély et al., 2016). Moreover, slow waves can emerge during wakefulness following sleep deprivation (SD), leading to the idea that sleep is an activity-dependent process (Alfonsa et al., 2023; Andrillon et al., 2021; Krueger et al., 2019; Thomas et al., 2020a; Vyazovskiy et al., 2011). The cellular and network counterpart of EEG slow waves is the alternation of neuronal activity and silence, often referred to as ON/OFF periods (Vyazovskiy et al., 2009). The incidence and duration of OFF-periods were shown to reflect sleep-wake history (Thomas et al., 2020b), and their experimental induction may lead to a decrease in electrophysiological and molecular signatures of sleep pressure (Driessen et al., 2026).

Sleep-dependent homeostatic processes have also been linked to cellular maintenance and macromolecule biosynthesis, both essential for a broad range of brain functions. For example, extended wakefulness is associated with widespread transcriptional and translational changes, including the altered expression of genes involved in synaptic plasticity, protein synthesis, and cellular stress pathways (Abdalla et al., 2026a; Lyons et al., 2023; Mackiewicz et al., 2007, 2008; Mignot, 2008; Vanrobaeys et al., 2023). Earlier studies have posited that wakefulness is associated with heightened neuronal activity, synaptic plasticity and neurotransmission in local neuronal circuits and the whole brain (Cirelli et al., 2004; Cirelli & Tononi, 2000), all functions requiring protein synthesis and changes in gene expression (Cirelli et al., 2004; Ma et al., 2023; Tononi & Cirelli, 2014).

Protein synthesis is a tightly regulated process, of which the Unfolded Protein Response (UPR) in the endoplasmic reticulum (ER) is an essential regulator (Walter & Ron, 2011). The UPR is an evolutionarily conserved signalling pathway activated by the accumulation of misfolded proteins in the ER lumen, observed in all species studied to date (Cirelli et al., 2004; Hetz et al., 2020; Maret et al., 2007; Naidoo et al., 2005, 2007; Walter & Ron, 2011). Activation of the UPR is initiated through three major signalling branches mediated by inositol requiring enzyme-1 (IRE1), PKR-like endoplasmic reticulum kinase (PERK), and activating transcription factor-6 (ATF6). These ER stress sensors are regulated in part through their association with the ER ATP-dependent chaperone binding immunoglobulin protein (BiP). During ER stress, altered interactions with BiP and unfolded proteins promotes activation of the UPR sensors, triggering signalling cascades that regulate protein synthesis, chaperone production and protein degradation. These responses initially act to restore ER proteostasis, whereas prolonged or unresolved ER stress can engage apoptotic pathways (H. P. Harding et al., 2009; Shen & Prywes, 2005; Walter & Ron, 2011; Ye et al., 2000) (Figure 1). Previous studies have shown that SD increases the expression of UPR-related genes in the brain of several vertebrate and invertebrate species (Abdalla et al., 2026b; Cirelli et al., 2004, 2005; Ford et al., 2024; Mongrain et al., 2025a; Naidoo et al., 2005, 2007; Terao et al., 2006), consistent with the view that neuronal activity during wakefulness imposes increased biosynthetic demand on neuronal networks. Although SD leads to changes in both cortical activity and ER stress, it is not known whether and to what extent the altered neuronal activity contributes to the observed molecular changes.

**Figure 1.**
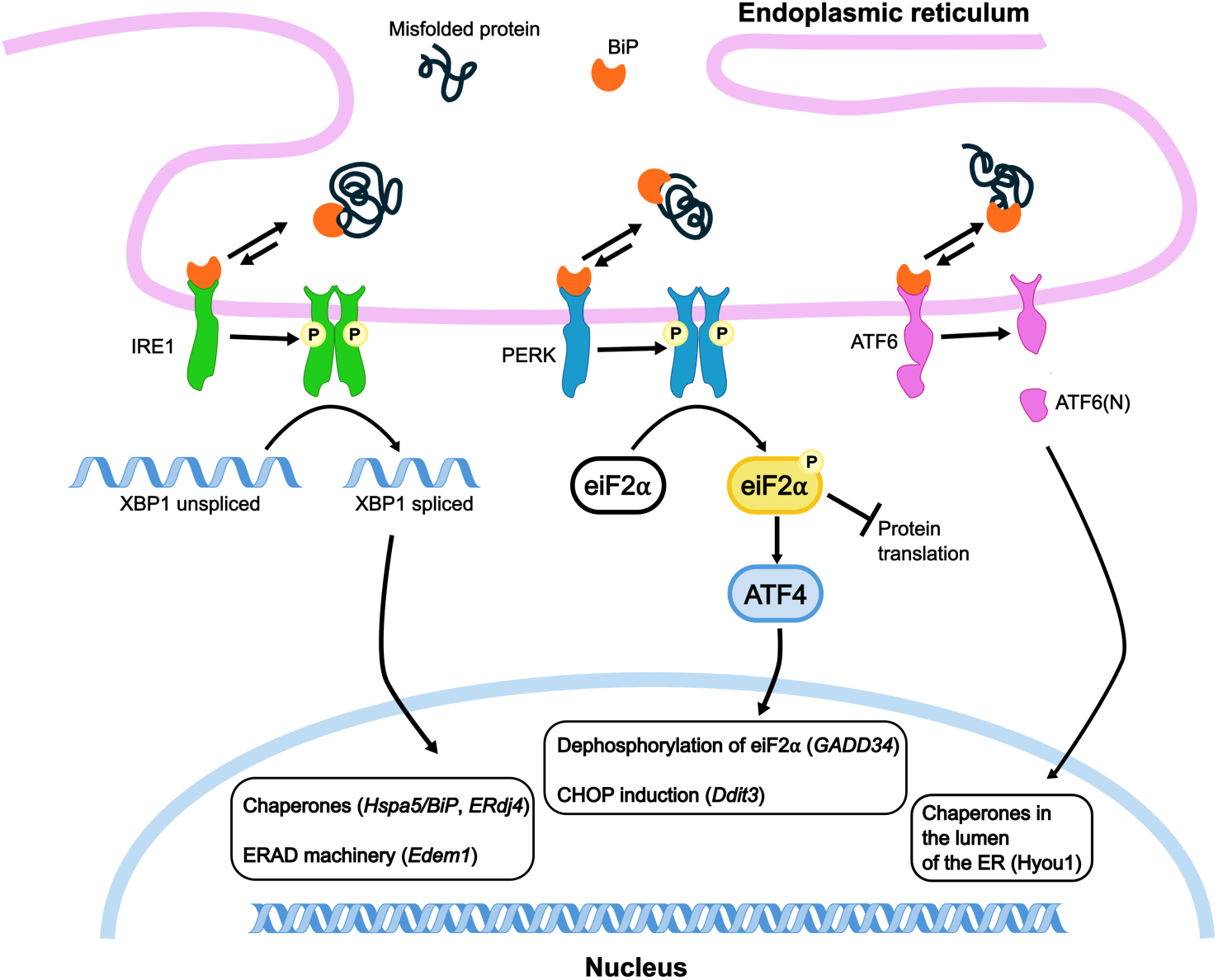
Core elements of the Unfolded Protein Response (UPR) signalling network. Misfolded proteins in the lumen of the ER activate the stress sensors IRE1, PERK and ATF6. Early response of the UPR generates active transcription factors to increase the protein-folding capacity of the ER while decreasing the load of proteins entering the ER. All three branches of the UPR are activated by dissociation of the ATP-chaperone protein BiP and its membrane receptor. Both IRE1 and PERK undergo phosphorylation and dimerization to reach an active configuration: IRE1 acts upon splicing transcription factor XBP1 to generate an active form capable of entering the nucleus, promoting the protein degradation machinery while increasing the synthesis of new chaperones. PERK phosphorylates eiF2α, which inhibits protein translation in the ER. If ER stress persists, eiF2α will induce transcription factor ATF4, which would lead to the activation of the cell apoptotic program. ATF6 is spliced, and the excised portion acts as a transcription factor that promotes chaperone production in the lumen of the ER.

Here, we tested how changes in cortical neuronal activity and prior sleep-wake history interact to regulate cortical gene expression and electrophysiological signatures of sleep homeostasis. To this end, we combined SD with monocular flickering light stimulation (FLS) at 8 Hz, a physiological sensory manipulation previously shown to modulate cortical activity in the visual cortex (Manning et al., 2007; Manning & Uhlrich, 2009; Uhlrich et al., 2005). Using electrophysiological recordings and cortical region-specific molecular analyses, we found that SD alone increased both activity-dependent and UPR-related gene expression together with SWA during subsequent sleep. In contrast, combining SD with FLS produced increased neuronal OFF periods during waking and further enhanced SWA during subsequent sleep, but prevented the widespread increase in both activity-dependent and UPR-related gene expression. Our findings identify physiological manipulation of neuronal activity as a potent modulator of ER-stress-associated gene expression in the cortex.

## Methods

### Experimental animals

Experiments were carried out in adult male C57BL/6 mice. We designed and performed three different series of experiments.

In the first experimental series, we recorded EEG and EMG data to evaluate regional and state-dependent differences in brain activity. Two cohorts of n=8 animals, (n=16 animals total) were implanted with EEG screws bilaterally in frontal and occipital cortices and nuchal EMG wires and were subjected to 4h sleep-deprivation (SD) with or without monocular flickering light stimulation (FLS). All these animals were subsequently used to collect tissue for molecular biology analysis. EEG and EMG signals of sufficiently high quality for subsequent analyses were acquired in n=12 animals.

In the second experimental series, we recorded local field potentials (LFPs) and multi-unit activity (MUA) from the visual cortex. Two cohorts of mice (n=4 and n=8, total of n=12 animals) were implanted with LFP electrodes unilaterally in the occipital cortex, EEG screws in contralateral occipital cortex and both frontal cortices, nuchal EMG and a head-mounted device for the FLS delivery to one eye. Good quality EEG, EMG, LFP and MUA signals were acquired in n=8 animals. A third cohort (n=4) of mice were implanted with EEG screws, LFP electrodes bilaterally in occipital cortex, nuchal EMG wires and had a head-mounted device for the monocular FLS device. In all cases, mice were subjected to 4h sleep-deprivation (SD) with and without FLS. For a subset of n=4, monocular FLS to the eye ipsilateral to the LFP electrode was performed instead of SD alone. The brains of these animals were subsequently processed histologically to confirm laminar probe location in cortex.

In the third experimental series, animals were intended exclusively for molecular biology analysis such as quantitative Polymerase Chain Reactions (qPCRs). It included a group of unimplanted animals (n=20 total) some of which were subjected to 4h SD (n=12) and some were left undisturbed for the duration of the experiment and served as controls (n=8). N=4 animals underwent surgery to attach the head-mounted monocular FLS device on top of the skull but had no EEG electrodes implanted. Further animals were EEG-screw implanted (but no EEG recordings were made) and either allowed to sleep (n=4; data not used) or underwent SD alone (n=12) or SD with FLS (n=4).

All animals used were between 7 and 10 weeks old at the start of the experiment. Mice were group housed until they underwent surgeries and singly housed subsequently. Following surgery, animals had a recovery period of at least seven days before being transferred into custom made Plexiglas cages (20.3 x 32 x 35 cm (W, D, H)) placed inside ventilated, sound attenuated Faraday cambers (Campden Instruments, Loughborough, UK) under standard 12h:12h light-dark cycle. Mice were habituated to the recording chambers and the tethering for 3 days before the start of recordings. Similarly, unimplanted mice for tissue collection were singly housed and habituated to the recording chambers for three days prior to the tissue collection day. Animals always had ad libitum access to food and water. Room temperature and relative humidity were maintained at 22±1°C and 50±20%, respectively. All procedures conformed to the Animal (Scientific Procedures) Act 1986 and were performed under UK Home Office Project and Personal Licenses in accordance with institutional guidelines.

### Surgical procedures

To study the effects of SD and FLS on brain activity, we performed surgical procedures for implantation of EEG screws, LFP electrodes, EMG wires and/or skull-mounted attachment of the screw for the monocular FLS device under aseptic conditions using isoflurane anaesthesia (3-5% for induction and 1-2% for maintenance). Three days before surgeries, mice started receiving plain jelly and mash daily for habituation to post-operative care. The day of the surgeries, Metacam (1–2 mg kg, subcutaneous (s.c)), and vetergesic (0.08 mg/kg, s.c.) were administered preoperatively. EEG/EMG mounts were composed of stainless-steel screws (shaft diameter,0.86 mm; InterFocus Ltd., UK) soldered to custom-made headmount connectors (8415-SM, PinnacleTechnology Inc., USA) and bilaterally implanted into frontal (motor area, anteroposterior (AP) 2 mm, mediolateral (ML) ±2 mm) and occipital cortical areas (visual area, V1, AP 3.5–4 mm, ML ±2.5 mm). A reference electrode was implanted above the cerebellum and two single stranded stainless-steel wires were inserted either side of the nuchal muscle to record electromyography (EMG) activity. All screws and wires were secured to the skull using dental cement (Super Bond Bulk mix kit, Sun Medical Co., LTD). The small attachment for the LED headstage was mounted during the surgery on the anterior portion of the acrylic head mount. To study neuronal responses to the FLS, laminar probes were implanted in one or both primary visual cortices (visual area, V1, AP 3.5 mm, ML ± 2.5 mm). All laminar recordings were performed using 16-channel silicon probes (NeuroNexus Technologies Inc., Ann Arbor, MI, USA; model: A1×16-3mm-100-703-Z16) with a spacing of 100 μm between individual channels. The tips of laminar implants were stained before surgery with the orange-red fluorescent membrane stain Dil ® (1,1′-Dioctadecyl-3,3,3′,3′-Tetramethylindocarbocyanine Perchlorate; Thermo Fisher Scientific, Waltham, MA, USA) by immersion of the electrode shank into a 20 mg/ml solution (in 50/50% acetone/methanol) for later histological assessment of the electrode position (Krone et al., 2021; Magill et al., 2006). After completion of the experiments, microlesions of selected channels on the laminar probe were performed under terminal pentobarbital anaesthesia using the electroplating device NanoZ™ (White Matter LLC, Seattle, WA, USA) applying 10 mA direct current for 25 s to each respective channel. Immediately following microlesioning, mice were perfused with 0.01M phosphate buffered saline (PBS; P4417, Sigma-Aldrich) followed by 4 % formaldehyde (F4775, Sigma-Aldrich) in PBS. A vibrating microtome (Leica VT1000S) was used to section the brains into 50 μm coronal slices. Slices in the appropriate region were stained with DAPI (4’,6-diamidino-2-phenylindole; D3571, Sigma-Aldrich, at 5μg/ml in PBS) at room temperature for one hour before being mounted and coverslipped. Slices were imaged using a Leica epifluorescence microscope (DMR) equipped with a Leica DC500 CCD camera. Images of implantation sites were aligned to the Paxinos and Franklin mouse brain atlas (Keith B.J. Franklin & George Paxinos, 2019) and the depth of the laminar implant was assessed measuring the distance between cortical surface and the electrical current induced tissue microlesions. The histological assessment of the insertion tract was performed in all animals, which were implanted with laminar probes.

### Experimental design

To enable a direct comparison of electrophysiological and molecular markers of neuronal activity and ER stress, we performed two types of complementary experiments. Firstly, to confirm regional variation in brain activity and assess the effect of monocular FLS, we used mice implanted with bilateral frontal and occipital EEG screws, a subset of which were also implanted with LFP electrodes. Secondly, to assess cellular activity at the molecular level and the unfolded protein response (UPR), brains of EEG implanted mice were used for regional tissue collection and qPCR after electrophysiological recordings, together with brains from unimplanted animals. Brain tissue samples were collected around ZT4 both for implanted and unimplanted animals, as well as for undisturbed and sleep-deprived mice. EEG implanted animals typically underwent chronic EEG recordings combined with SD and/or SD+FLS on 1-3 occasions prior to the day of tissue collection (n=16), except for two groups (n=12) that received single SD+FLS on the day of tissue collection and were not used for chronic EEG recordings.

Animals were continuously recorded for EEGs (n=16) and LFPs (n=8), starting at ZT0 when the welfare checks were also performed. On SD days with or without monocular FLS stimulation, mice were subjected to 4 hours of SD through the presentation of novel objects from light onset under continuous monitoring (ZT0-ZT4), as previously described (McKillop et al., 2018; Yamagata et al., 2021a). FLS consisted of 8Hz trains of white light emitted from a small LED device, lasting for 2 seconds and repeated every 30s during the entire duration of the 4-h SD. LED intensity was optimised to reliably induce visually detectable cortical responses at a single trial level, and maintained constant during the stimulation. The temporal frequency was chosen based on previous work in rats (Manning et al., 2007; Uhlrich et al., 2005) where it was shown to elicit the strongest evoked EEG response in the visual cortex and potentiation of responses across repeated sessions. Stimulation was presented using custom-made headstages containing one light-emitting diode (Luxeon Star LED, product code: LXML-PWC2 5650K) at a viewing angle of 30 degrees, 6-8 mm from the corneal surface of the eye, positioned over the centre of the pupil. The LED device was attached at the beginning of the 4-h SD period and removed immediately before the animals were left undisturbed. Stimuli were triggered externally by the computer, using the software and hardware from Tucker Davis Technologies (11930 Research Circle Alachua, FL 32615 USA), which allowed precise synchronisation of EEG signals and LED stimulation for subsequent analysis.

### Signal processing and analysis

Electrophysiological recordings were acquired with an RZ2 High-Performance Processor and Synapse software (Tucker-Davis Technologies Inc., Alachua, FL, USA). EEG, EMG, and LFP signals were continuously recorded, concomitantly with extracellular neuronal spike data from the same electrodes used for LFP monitoring (PZ5 NeuroDigitizer preamplifier, TDT, USA). EEG, LFP, and EMG signals were filtered between 0.1 and 100 Hz and stored at a sampling rate of 305 Hz. The signals were resampled offline at a sampling rate of 256 Hz using custom-made Matlab scripts (The MathWorks Inc., Natick, MA, USA). For subsequent analyses, EEG and LFP power spectra were computed by a fast Fourier transform of 4-s epochs (Hanning window), with a 0.25-Hz resolution in Matlab. Signals between 0.5-20 Hz were analysed as changes in this range have been most consistently related to sleep pressure. Extracellular neuronal activity was continuously acquired at a sampling rate of 25 kHz and filtered between 300 Hz and 5 kHz. For spike acquisition, amplitude thresholds were manually set on Synapse on each recording channel. This threshold was set at least 2 standard deviations above noise level using automatic thresholding tool from Synapse that was manually reviewed afterwards. When the voltage crossed this predefined set threshold, 46 samples around the event (0.48 ms before and 1.36 ms after the threshold crossing) were extracted and recorded. Spike waveforms were processed using custom-made Matlab scripts. To screen out putative units of good quality for the 12-h light phase data of each channel, K-means analysis was performed using principal components from PCA analysis based on waveform data recorded via thresholding. Clustered waveforms were then visually inspected and screened. Subsequently, firing rate of each day were plotted and 6 std cutoff (or in two cases, 1 or 3 std) from the mean was applied for noise rejection. OFF periods were detected in each channel (selected best quality 2-3 channels per animal) as neuronal silence lasting at least 50 ms.

### Scoring and analysis of vigilance states

Sleep scoring was performed offline. Resampled EEG, LFP and EMG signals (256 Hz) were transformed into European Data Format (EDF) using custom made scripts in python (Python 3.10). These signals were visualized in 4-s epochs using the software SleepSign (Kissei Comtec Co., Nagano, Japan) and manually scored for vigilance state. Wake was defined as low-voltage, high-frequency EEG activity accompanied by a high level of EMG activity lasting more than 4 epochs. NREM sleep was defined as EEG signal with high voltage and slow frequency, predominantly characterized by the occurrence of slow waves (0.5 to 4 Hz) and sleep spindles (10-15 Hz), and low EMG activity. REM sleep was defined as low-voltage, high-frequency EEG oscillations, with predominance of theta (6 to 9 Hz) activity in the occipital derivations, which was distinguished from waking by reduced level of EMG activity. Brief awakenings (microarousals) were defined as transient periods of low-voltage, high-frequency oscillations in the EEG accompanied by elevated EMG tone, lasting ≥2 s and ≤16 s. Epochs containing artefacts, resulting from eating, drinking or gross movements were excluded from spectral/MUA analysis.

### RT-PCR

For RNA analyses, undisturbed (BL), sleep-deprived (SD) or sleep-deprived with monocular flickering light stimulation (SD+FLS) mice were sacrificed by cervical dislocation at ZT4 (see Methods Fig 1 for summary of experimental groups). Brains were rapidly removed and placed into a brain matrix (Kent Scientific). A skin graft blade (Swann-Morton) was used to dissect 1 mm thick brain slices. Tissue punches were taken from frontal cortex and visual cortex using a sample corer (1 mm internal diameter, Fine Science Tools GmBH), flash frozen on dry ice and stored at -80°C until use. For tissue RNA extraction, the tissue was firstly mechanically disrupted in 100 µl of Trizol. The sample was then made up to 500 µl with Trizol, 100 µl chloroform added and then thoroughly mixed. Following a 5 min incubation at RT, the sample was then centrifuged for 15 min at 15,000 xg, 4°C. The clear top layer was then carefully collected, mixed with an equal volume of 70% ethanol, and RNA extracted using the RNeasy Plus Mini Kit (Qiagen) following the manufacturer’s instructions. RNA concentration and quality was determined using a NanoDrop ND-1000 spectrophotometer and cDNA synthesized using the qScript cDNA synthesis kit (QuantaBio). RT-PCR was then conducted using the Quantifast SYBR Green PCR Kit (Qiagen) and a StepOnePlus thermal cycler (Applied biosystems) with the following thermal profile: 95°C for 5 mins and then 40 cycles of 95°C for 10s, 60°C for 30s and 72°C for 12 s. Quantification of transcript levels was conducted using the relative standard curve method, for comparing within a gene, or the 2-ΔCt method for comparison across genes or samples, using GAPDH and RPL19 as housekeeping genes for the normalisation.

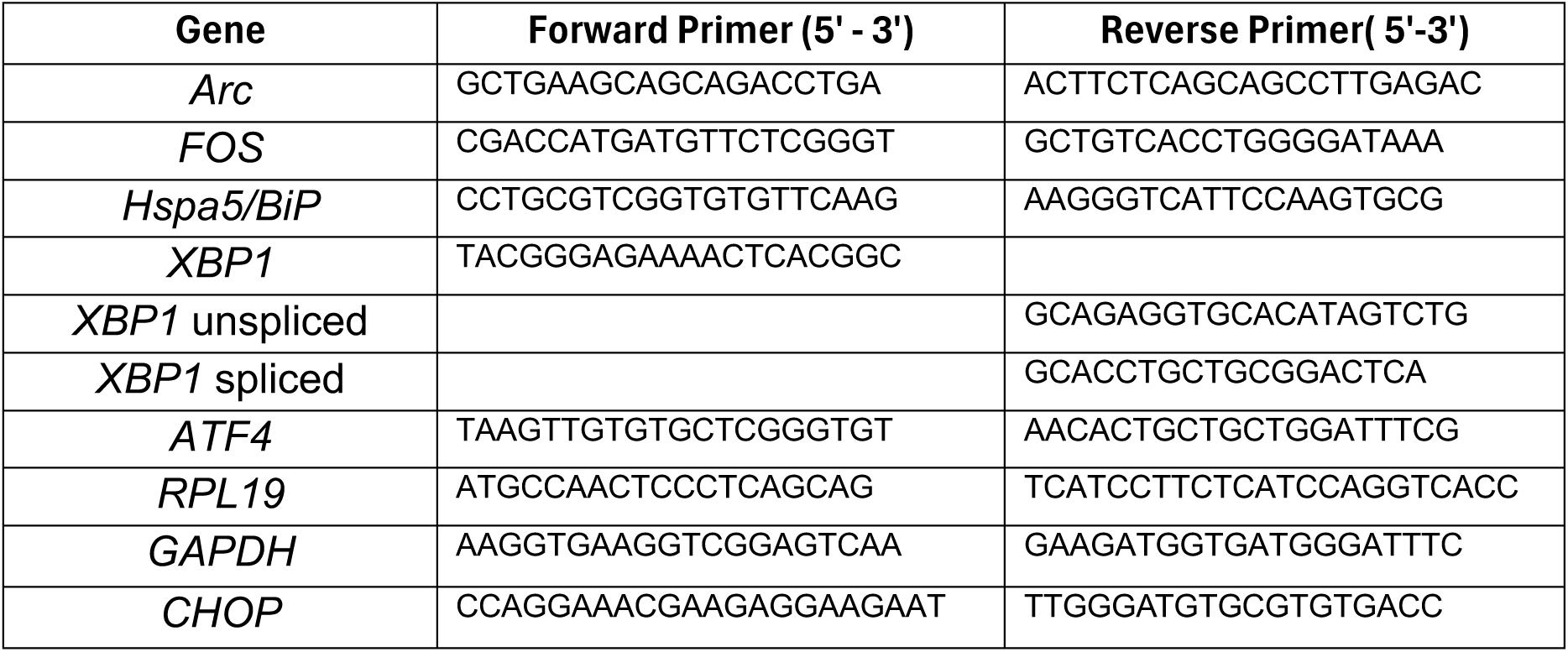

### Statistics

GraphPad Prism Version 10.4.1 was used for all statistical analysis. For RT-PCR analysis, Analysis of Variance for comparison of different experimental conditions were performed using two-way ANOVA with post-hoc tests corrected for multiple comparisons with Šídák multiple comparison test, alpha 0.05 for SD vs baseline (n=8 mice for baseline and n=11 mice for SD). Similarly, for SD+FLS vs SD comparison, Welch’s t-test was used (n=16 mice for SD and n=16 or 32 mice for SD+FLS). For both absolute and relative power density spectrum analysis of EEG recordings, two-way ANOVA with Šídák’s multiple comparisons test with factors “frequency bins” and “derivations” or Wilcoxon matched-paired signed rank test were used, alpha 0.05. Matlab was used for statistical analysis of MUAs. Statistical comparisons were performed using a two-tailed paired t-test was performed with units from best quality selected channels in the visual cortex, n=14 channels from n=5 mice, each with SD and SD+FLS conditions as well as respective baseline days.

## Results

### Sleep deprivation-induced changes in cortical slow-wave activity and expression of neuronal activity- and UPR-related genes

To evaluate the relationship between cortical activity, sleep-wake dynamics and gene expression, mice were implanted with frontal and occipital EEG electrodes and recorded during undisturbed 24-h baseline and SD (Fig. 2A). Baseline recordings under 12h-12h light-dark cycle allowed the determination of vigilance states (Fig. 2B) and showed that mice spent a high proportion of time asleep during the light period and awake during the dark phase. As expected, slow-wave activity (SWA; 0.5–4 Hz) during NREM sleep was greater in the frontal than in the occipital derivation, whereas the opposite pattern was observed for theta activity during wakefulness and REM sleep (Fig. 2C). Mice were recorded continuously, with SD performed from ZT0 to ZT4 following 24 h of baseline recording (Fig 2D). Compared to spontaneous wakefulness, SD resulted in increased EEG power in both SWA and theta ranges in both derivations, with the increase at SWA being more pronounced in the frontal cortex. During subsequent NREM sleep, consistent with previous findings (Guillaumin et al., 2018), the increase in SWA was greater in the frontal than in the occipital derivation (Fig. 2E). For molecular assays, a separate cohort of animals were perfused at ZT4, with or without 4h of preceding SD. Unlike its more pronounced effect upon frontal EEG, the effect of SD upon gene expression tended to be comparable across cortical regions, or were more pronounced in occipital cortex (Suppl. Fig. 1). In the frontal derivation, SD resulted in a 2-fold or higher increase in both *Fos* and *Arc* expression, although high variability among animals in *Fos* expression was noted (*Arc*: p=0.0001; *Fos*: p=0.006; Holm-Šídák’s multiple testing correction). SD alone also increased by at least 1.5-fold the expression of multiple UPR-related genes in the frontal cortex, including *Hspa5/BiP*, *Xbp1*-spliced and unspliced, *Gadd34* and *Hyou1* (*Hspa5/BiP*: p<0.0001; *Xbp1*-spliced: p<0.0001; *Xbp1*-unspliced: p=0.02; *Hyou1*: p<0.0001; *Gadd34*: p<0.0008; Holm-Šídák’s multiple testing correction). In contrast, the expression of genes associated with later-stage UPR signalling and pro-apoptotic pathways, including *Atf4* and *Ddit3*/*Chop*, was not altered by SD alone in the frontal derivation. In the occipital cortex, SD resulted in at least a 2-fold increase in both *Fos* and *Arc* levels (*Arc*: p<0.0001 and *Fos*: p=0.015; Holm-Šídák’s multiple testing correction). Likewise, expression of *Hspa5/BiP*, *Xbp1*-spliced and unspliced, and *Hyou1* was increased at least 2-fold following SD (*Hspa5/BiP*: p<0.0001; *Xbp1*-spliced: p=0.003; *Xbp1*-unspliced: p=0.0006; *Hyou1*: p=0.0008; Holm-Šídák’s’s multiple testing correction), while Gadd34 was still increased approximately 1.4-fold (*Gadd34*: p=0.021; Holm-Šídák’s’s multiple testing correction) (Fig. 2F). Because both forms of the XBP1 transcript are increased during SD, it may reflect increased XBP1 transcript abundance in addition to IRE-1 dependent splicing.

**Figure 2.**
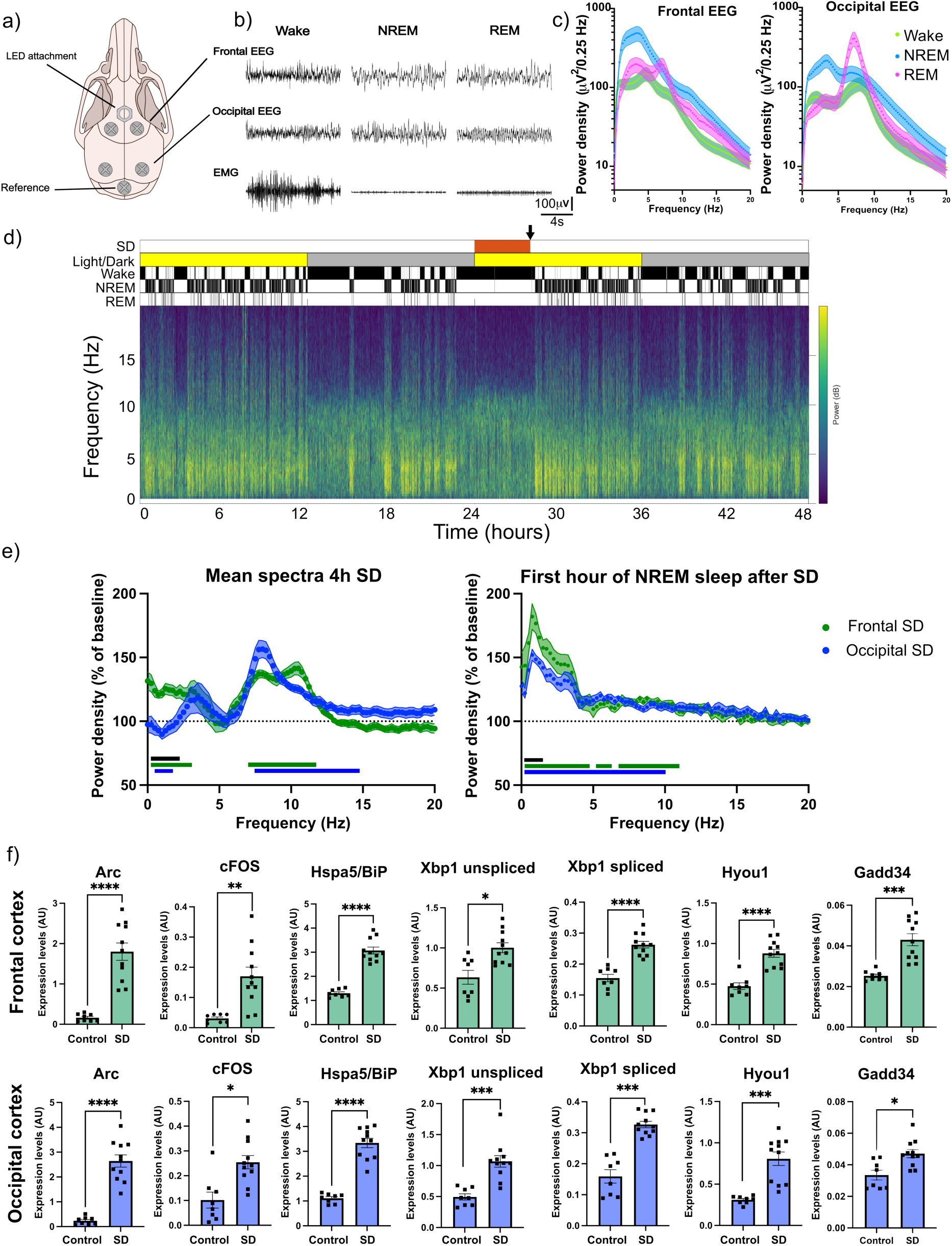
Sleep deprivation increases electrophysiological signatures of sleep pressure and upregulates activity-dependent and ER-stress related genes in frontal and occipital cortex. **A)** Configuration of EEGs and attachment for LED light on the mouse skull. **B)** Traces of EEGs and EMGs during wake, NREM sleep and REM sleep in a representative animal. **C)** Average EEG power spectra during 24-h baseline recordings (n=8) shown for each vigilance state in the frontal and occipital EEGs. **D)** Representative 48-h frontal EEG spectrogram (bottom) aligned with the time where sleep deprivation was performed (red) and a vertical arrow showing the average point of tissue collection for RT-PCR analysis, the 12h-12h light/dark cycle, and the hypnogram showing wake, NREM and REM. **E) Left** panel shows EEG power density for frontal (green) and occipital (blue) signals during the entire 4-h sleep deprivation, expressed as percentage of the same frequency bin for wake epochs during the 24-h baseline (mean ± S.E.M.). Two-way ANOVA with Šídák’s multiple comparisons test with factors “frequency bins” and “derivations”; Frequency - F (120, 1650) = 15.81, p<0.0001, Derivation - F (1, 1650) = 78.50, p<0.0001, Interaction - F (120, 1650) = 4.078, p<0.0001. **Right** panel shows power density of NREM epochs for frontal (green) and occipital (blue) EEG signals during the first hour after sleep deprivation, expressed as percentage of the same frequency bin for 24-h baseline NREM epochs (mean ± S.E.M.). Two-way ANOVA with Šídák’s multiple comparisons test with factors “frequency bins” and “derivations”; Frequency - F (120, 1642) = 25.58, p<0.0001, Derivation - F (1, 1642) = 13.60, p=0.0002, Interaction: F (120, 1642) = 1.560, p=0.0002. Black horizontal bars denote frequency bins where the difference between the frontal and occipital derivation was significant (p<0.01). Green and blue horizontal bars indicate significant differences in mean power relative to baseline (p < 0.01), using Wilcoxon matched-paired signed rank test without corrections for multiple comparisons, n=8. **F)** Effect of sleep deprivation on gene expression in frontal cortex (top, green bars) and occipital cortex (bottom, blue bars). In frontal cortex, activity-dependent genes (*Arc* and *Fos*) and multiple ER-stress related genes are significantly upregulated after SD (*Arc*: p=0.0001; *cFos*: p=0.006; *Hspa5*/BiP: p<0.0001; *Xbp1*-spliced: p<0.0001; *Xbp1*-unspliced: p=0.02; *Hyou1*: p<0.0001; *Gadd34*: p<0.0008; Welch’s t-test using Holm-Šídák’s multiple testing correction). Similarly, in occipital cortex, activity-dependent genes and multiple ER-stress related genes are significantly upregulated (*Arc*: p<0.0001 and *Fos*: p=0.015; Holm-Šídák’s multiple testing correction; *Hspa5*/BiP: p<0.0001; *Xbp1*-spliced: p=0.003; *Xbp1*-unspliced: p=0.0006; *Hyou1*: p=0.0008; *Gadd34*: p=0.021; Holm-Šídák’s’s multiple testing correction, control n=8, SD n=11).

To further explore whether expression of ER-stress-related genes was associated with neuronal activity, we performed correlation analyses between activity-dependent genes (i.e., *Fos* and *Arc*) and UPR-related transcripts. Overall, the analyses revealed positive associations, with correlations between *Arc* and UPR-related genes generally stronger than those involving *Fos*, although rested and sleep-deprived animals formed largely distinct clusters (Supplementary Fig. 2).

### Flickering light stimulation induces neuronal OFF periods during sleep deprivation and leads to an increase in SWA during subsequent sleep

As SD leads to global changes in electrophysiological and molecular markers of neuronal activity, we hypothesised that the changes in cortical activity during SD are responsible for the molecular signatures associated with ER stress. To address this question, we combined SD with monocular FLS in a separate cohort of mice. The stimulation paradigm consisted of pulses of bright white light delivered to one eye via a skull-mounted LED, which allowed the animal to move freely throughout the stimulation (Figure 3A). Trains of 10-ms light pulses at 8 Hz were delivered for 2 s every 30 s throughout the 4-h SD period, and reliably affected activity in the contralateral primary visual cortex as recorded by intracranial laminar probe (Figure 3B). We chose the 8 Hz frequency to further enhance the spontaneous oscillatory activity typical of active exploratory behaviour and high arousal levels (Yamagata et al., 2021b).

**Figure 3.**
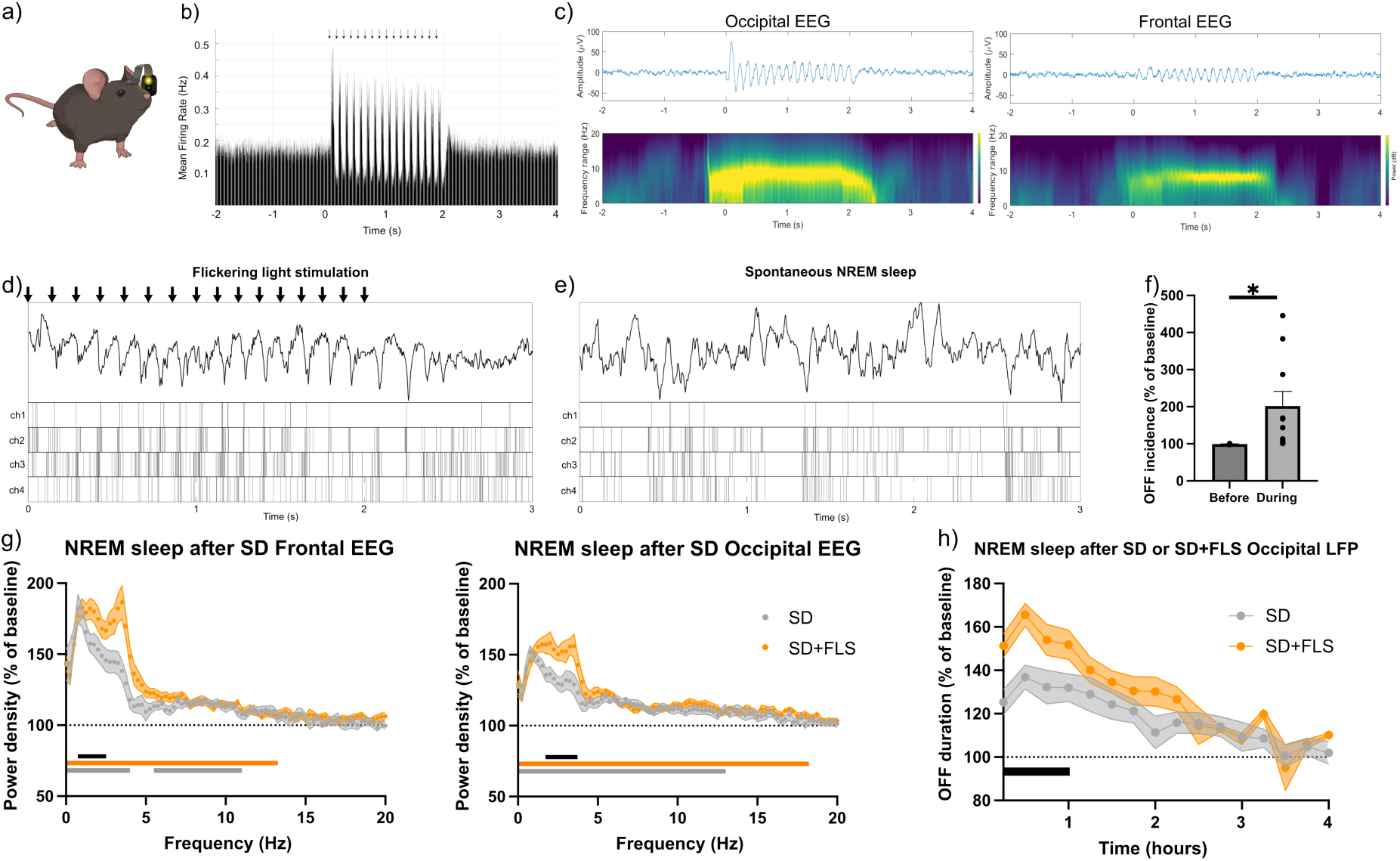
Effects of flickering light stimulation upon cortical neuronal activity and EEG during waking and subsequent sleep. **A)** Schematic showing the LED device for delivering flickering light stimulation (FLS). **B)** Average multiunit activity (MUA) during s epochs across multiple trains (n=480 total) of FLS (8 Hz for 2s, 16 pulses total) in a representative animal. **C)** Mean EEG traces and spectrogram (n=480 traces per animal) around the onset of FLS for frontal and occipital EEGs in a representative animal. **D)** Example LFP trace and MUA during a single train of FLS. **E)** Example LFP trace and MUA during representative 3s of spontaneous NREM sleep. In both D) and E) the raster plots below LFP traces represent the occurrence of spikes in individual channels of the laminar probe. **F)** Comparison of OFF period incidence for 2s before and during the 2s FLS train, mean±SEM (before stimulation: 99.28±0.32; during stimulation: 201.9 ±39.74 (One-tailed t-test, p=0.0147, alpha=0.05)). All values are relative to baseline OFF incidence (n=10 LFP channels with good MUA from n=5 mice). **G)** Frontal (left) and occipital (right) relative EEG power density spectra for the first hour after SD or SD+FLS (mean ± S.E.M.). Two-way ANOVA with Šídák’s multiple comparisons test with factors “frequency bins” and “experiment”; Frontal: Frequency - F (120, 2142) = 42.78, p<0.0001, experiment - F (1, 2142) = 58.74), p<0.0001, interaction - F (120, 2142) = 2.482, p<0.0001. Occipital: Frequency - F (120, 2142) = 29.11, p<0.0001, experiment - F (1, 2142) = 56.91, p<0.0001, interaction: F (120, 2142) = 1.603, p<0.0001. **H)** Comparison of the relative OFF period duration in artifact-free NREM epochs during 4 hours after SD or SD+FLS, expressed as percentage of baseline NREM OFF period duration. It was analysed using a two-way repeated-measures ANOVA for each hour interval, with experimental condition (SD and SD+FLS) and time (4 consecutive 15-minute intervals) as within-subject factors. The Geisser– Greenhouse correction was applied, and post hoc comparisons between conditions at individual time points were adjusted for multiple testing, showing significant differences, represented by black lines, only during the first hour after SD or SD+FLS; Time Interval: F (3, 24) = 5.936, p=0.0035; Condition: F (1, 8) = 8.952, p=0.0173; Interaction: F (3, 24) = 0.8221, p=0.4945.

Directly observing the animals confirmed that they tolerated the stimulation paradigm well and did not manifest obvious alterations in spontaneous behaviour. Furthermore, the total amount of waking during SD was not significantly affected by the procedure (3.57±0.11 vs 3.64±0.08 hours for SD vs SD+FLS respectively, unpaired two-tailed Welch’s t-test, p=0.6). As the animals were handled to disconnect the device at the end of the SD+FLS procedure, they took slightly longer to fall asleep, as compared to the SD alone condition (NREM during the 1st hour after SD and SD+FLS was 0.37±0.04 and 0.22±0.05 hours respectively, unpaired two-tailed Welch’s t-test, p<0.05).

As expected, FLS elicited a strong theta (8Hz) oscillation in the EEG and LFP in the visual cortex, which was present, although in a somewhat attenuated form, also in the frontal derivation (Figure 3C). Overall average firing rate during wake was lower in the SD+FLS condition as compared to SD alone (SD alone = 138.1 ± 12.1%; SD+FLS = 125±9.9%, as percentage of baseline; p=0.043 two-tailed paired t-test). This change in average firing rate was explained by the markedly altered firing pattern during stimulation. Neuronal activity recorded from the visual cortex was strongly entrained by the FLS, showing brief bursts of activity, that were followed by a prominent suppression in neuronal spiking (Figure 3D). These suppressions in MUA resembled OFF periods typically observed during sleep (Figure 3E) (Harding et al., 2023), and a quantification of OFF periods revealed that their incidence doubled during delivery of the FLS compared with the immediately preceding period (Figure 3F).

Despite the increase in the incidence of sleep-like OFF periods induced by stimulation, we did not detect a significant difference in subsequent sleep duration of animals subjected to light stimulation during SD, when compared to those subjected to SD alone (ZT4-ZT24, 10±0.38 hours (n=8) vs 8.78±0.37 hours (n=12) for SD alone or SD+FLS respectively, paired two-tailed Wilcoxon test, p=0.0781). However, the subsequent sleep was characterised by higher EEG SWA in both the occipital and frontal derivations (Figure 3G), as well as longer duration of OFF periods (Figure 3H). This suggests that there was a global increase in sleep pressure in response to physiological stimulation, even if it was associated with the induction of local OFF periods.

### Flickering light stimulation abolishes sleep-deprivation-induced changes in gene expression

To investigate whether local stimulation led to changes in neuronal activity-related and ER stress-related genes, we repeated the transcriptomic analysis on cortical tissue punches collected immediately after SD+FLS and compared these results to the SD only condition. Gene expression in the hemisphere contralateral to the eye that received stimulation was compared with the averaged gene expression across both hemispheres in SD only brains. Remarkably, the increases in activity-related genes due to SD were no longer observed in visual cortex. Both *Fos* and *Arc* showed a marked decrease (1.96-fold for *Fos* and 3.7-fold for *Arc;* p<0.0001 for both genes, Holm-Šídák’s’s multiple testing correction) in the SD+FLS condition when compared to SD alone (Figure 4A). The expression of genes encoding members of ER stress pathways upregulated by SD was also attenuated following FLS*. Hspa5*/BiP, *Xbp1*-spliced and unspliced, and *Hyou1* were all reduced in the SD+FLS condition by at least 1.8-fold (p<0.0001 for all four genes; Holm-Šídák’s’s multiple testing correction; Figure 4B). Expression of *Hspa5*/Bip, which acts as a main sensor of UPR pathways during ER stress, was also attenuated (2.90±0.91 vs 1.45±0.48 for SD alone vs SD+FLS respectively; expression levels: AU, mean±s.d.). Spliced *XBP1* (*XBP1s*), which encodes a potent transcriptional activator, and unspliced XBP1 (*XBP1u)*, which encodes an inhibitor of the UPR (Yoshida et al., 2006), were both attenuated (*XBP1s* 0.3±0.06 vs 0.16±0.05, and *XBP1u* 0.93±0.35 vs 0.5±0.14, for SD alone vs SD+FLS, respectively) suggesting that FLS primarily attenuated the SD-associated increase in XBP1 transcription rather than selectively altering the balance between the spliced and unspliced isoforms. Meanwhile, the expression of *Hyou1*, a chaperone with a similar function to *BiP* (Kusaczuk & Cechowska-Pasko, 2013) was also attenuated (0.70±0.30 vs 0.33±0.07 for SD alone vs SD+FLS respectively). These population data were pooled from three different experimental batches and consistent effects were observed across each of the three batches.

**Figure 4.**
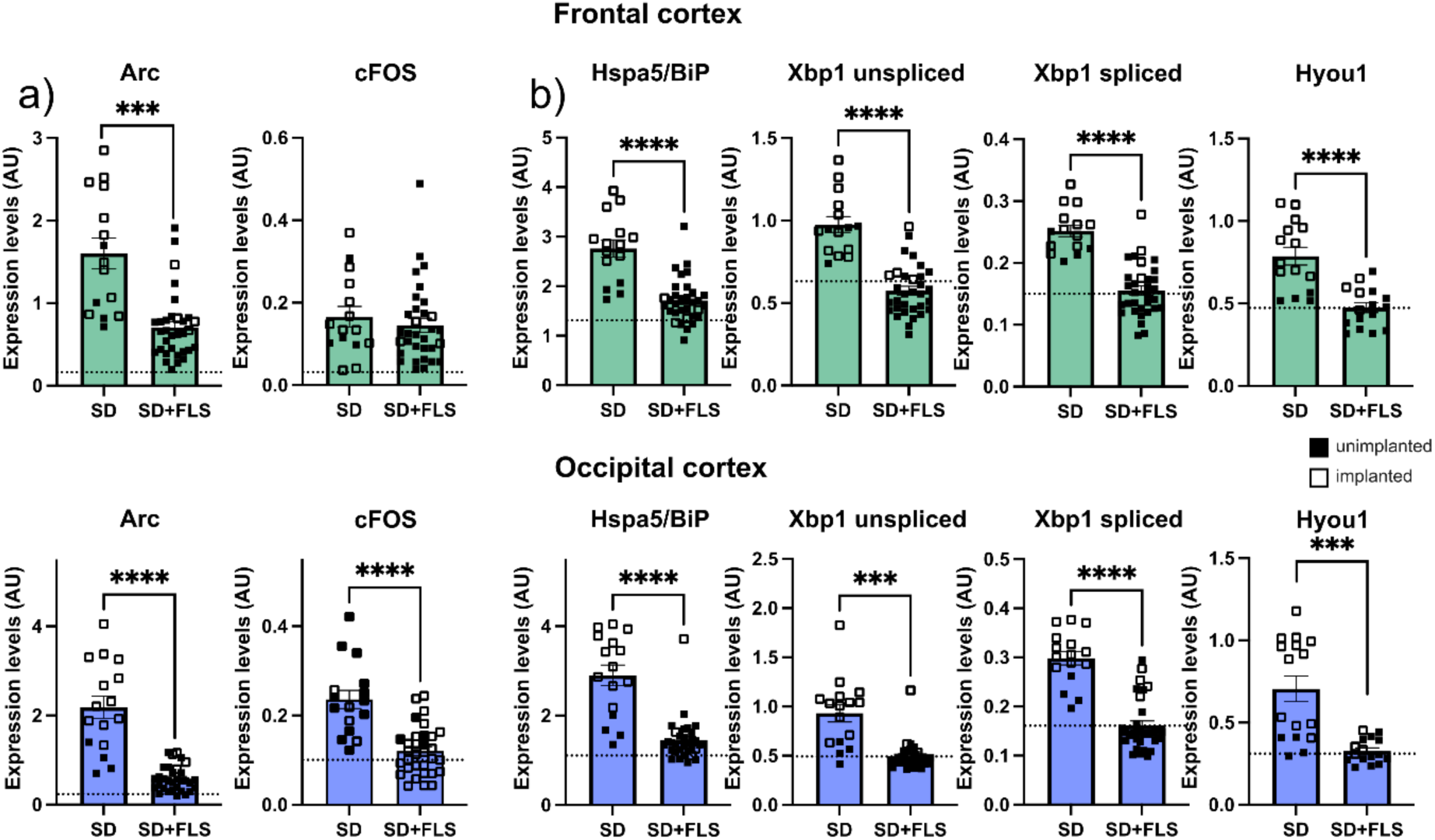
FLS during sleep deprivation prevents increases in activity-dependent and ER-stress related genes in different cortical regions. **A)** Gene expression levels of the immediate early genes *Fos* and *Arc* as measured by qPCR in frontal and occipital cortices of mice that were either sleep-deprived (SD) or sleep-deprived with flickering light stimulation (SD+FLS) for 4hrs. Tissue samples were taken from the hemisphere contralateral to the eye receiving the stimulation in the preceding 4h period. Expression of *Arc* is significantly reduced by FLS in frontal cortex, while both genes are downregulated in occipital regions (Frontal cortex: *Arc*: p<0.0001; *Fos*: p>0.9999; Occipital cortex; p<0.0001 for both genes, Holm-Šídák’s’s multiple testing correction). **B)** Gene expression levels of members of the ER-stress pathway. Three genes were significantly downregulated following FLS in both frontal and occipital cortices (at least p<0.001 for all four genes; Holm-Šídák’s multiple testing correction). *p<0.05, ** p<0.01, *** p<0.001, **** p<0.0001. SD is n=15 of which 4 are implanted and 11 are unimplanted and SD+FLS is n=32 of which 28 are implanted and 4 are non-implanted. Dashed line represents the expression of the gene presented in undisturbed condition, when animals are allowed to sleep at will. Solid symbols represent mice not implanted with EEG headstages.

A similar profile was observed in frontal cortical regions, where the FLS attenuated upregulation of SD-induced genes, with the exception of *Fos*. Specifically, the activity regulated gene *Arc* was expressed at 2-fold lower levels when FLS was delivered compared to SD alone (1.60±0.72 vs 0.71±0.41 for SD vs SD+FLS, p=0.0003 Holm-Šídák’s’s multiple testing correction). Similarly, *Hspa5/*BiP, *XBP1u*, *XBP1s,* and *Hyou1* were all approximately 1.6-fold reduced after SD+FLS compared to SD alone (*Hspa5*: 2.76±0.68 vs 1.71±0.45, p<0.0001; *XBP1u*: 0.98±0.19 vs 0.58±0.15 p<0.0001; *XBP1s*: 0.25±0.04 vs 0.16±0.04 p<0.0001; *Hyou1*: 0.79±0.21 vs 0.47±0.12 p=0.0002; mean±s.d. and Welch’s t-test using Holm-Šídák’s’s multiple testing correction).

## Discussion

Previous work has established that prolonged wakefulness is accompanied by increased neuronal activity, elevated expression of immediate early genes, and activation of multiple branches of the UPR (Abdalla et al., 2026c; Basheer et al., 1997; Cirelli et al., 2004; Mackiewicz et al., 2007; Mongrain et al., 2025b; Naidoo et al., 2005, 2007; Popescu et al., 2025). These observations have been interpreted as reflecting the increasing biosynthetic demands imposed on cortical neurons during sustained wakefulness (Vanrobaeys et al., 2023; Vyazovskiy & Harris, 2013). We reveal that although electrophysiological markers of sleep pressure (recovery sleep SWA) exhibit substantial heterogeneity across cortical regions, the molecular response to prolonged wakefulness appears to be more uniform across the cortex. This observation suggests that cortical activity underlying transcriptional regulation is not simply reflected by the spatial distribution of SWA.

When neuronal activity was manipulated during sleep deprivation by flickering light stimulation, we observed transient suppressions in neuronal firing, while the total duration of wakefulness was unchanged. Critically, however, manipulating neuronal activity in this manner prevented the increase in activity-dependent and UPR-related gene expression induced by sleep deprivation. These results suggest that the UPR is influenced not simply by the duration of prior wakefulness, but by the temporal pattern or type of neuronal activity during waking. Interestingly, reduction of UPR-related gene expression during SD occurred despite a further increase in subsequent SWA. To our knowledge, this is the first experimental demonstration that molecular markers of ER stress and electrophysiological markers of sleep homeostasis can be uncoupled through physiological manipulation of cortical neuronal activity. While SWA remains one of the most robust measures of accumulated sleep pressure, our findings suggest that it does not fully capture the molecular state of cortical neurons. Instead, different components of neuronal activity may regulate distinct aspects of the response to prolonged wakefulness.

One possible interpretation of the differential effects of neuronal activity changes in markers of sleep pressure such as SWA and ER-stress related gene expression could be that the UPR is primarily sensitive to action potential firing and the associated metabolic and biosynthetic demands, whereas SWA may be more strongly influenced by synaptic and subthreshold network activity. Repeated firing generates large intracellular calcium transients, increases vesicle recycling, membrane trafficking, and protein synthesis, all processes that place considerable demands on endoplasmic reticulum proteostasis (Brini et al., 2014; Cohen & Greenberg, 2008; Ma et al., 2023; Rangaraju et al., 2017; Walter & Ron, 2011). By repeatedly interrupting neuronal firing through induced sleep-like OFF periods, FLS may reduce these biosynthetic demands sufficiently to attenuate activation of UPR pathways. At the same time, repeated stimulation may favour the recruitment of cortical inhibitory neurons such as somatostatin positive neurons and contribute to the accumulation of sleep pressure (Funk et al., 2017), explaining the enhanced SWA observed during subsequent recovery sleep. Although this model remains speculative, it provides an explanation for the apparent dissociation between transcriptional and electrophysiological markers of sleep pressure observed in the present study. This interpretation is also consistent with recent work showing that experimentally induced OFF periods promote synaptic renormalisation and weaken excitatory cortical synapses (Driessen et al., 2026). Our findings extend these observations by suggesting that OFF periods may not only influence synaptic physiology but also regulate activity-dependent transcriptional programmes associated with ER stress.

The broader implication is that distinct features of cortical activity may regulate different biological processes. We speculate that sustained action potential firing may primarily determine activity-dependent transcription and engagement of protein quality-control pathways, whereas other forms of neuronal activity, including synaptic and subthreshold signalling, may contribute more strongly to the accumulation of sleep pressure reflected by SWA. Future experiments combining intracellular recordings, calcium imaging, and direct measurements of protein synthesis and ER function will be required to test this hypothesis.

Finally, the findings of the current study may have implications beyond sleep physiology. Dysregulation of ER proteostasis and UPR signalling has been implicated in several neurological disorders, including Alzheimer’s disease and epilepsy. Notably, non-invasive sensory stimulation has emerged as a potential strategy for modifying disease-associated cellular processes. 40-Hz visual stimulation reduces amyloid-β burden and alters microglial responses in mouse models of Alzheimer’s disease, while prolonged or multisensory gamma entrainment influences neuronal and synaptic integrity, glial responses, and cognition (Adaikkan et al., 2019; Adaikkan & Tsai, 2020; Iaccarino et al., 2016; Martorell et al., 2019). Although our stimulation paradigm differs in frequency and experimental context, our findings raise the possibility that sensory stimulation can additionally influence cellular stress pathways through activity-dependent mechanisms. Determining whether modulation of ER proteostasis contributes to the beneficial effects reported for sensory stimulation in disease models will be an important direction for future studies.

## Author contributions

Conceptualization and design: J.P.M, V.V.V.; Methodology: J.P.M., L.T., A.H.S.; Investigation: J.P.M., L.T., A.H.S., A.C.; Funding acquisition: V.V.V, C.J.A.; Project administration: J.P.M., V.V.V.; Supervision: V.V.V., C.J.A.; Writing, original draft: J.P.M.; Writing, review and editing: J.P.M., S.E.N., C.J.A., V.V.V., A.J., R.M., A.H.S.

## Acknowledgments

We thank the members of Aarti Jagannath’s, Stuart Pierson’s and Colin J. Akerman’s labs for helpful discussions. We thank Lukas Krone, Cristina Blanco-Duque, Martin Khan, Linus Milinski, Sian Wilcox and Laura McKillop from Vladyslav Vyazovskiy’s lab for their assistance during the *in vivo* experiments and guidance.

## Funding

This research received funding from the Welcome Trust Grant 227093/Z/23/Z, Medical Research Council (MRC) projects MR/S01134X/1 and MR/Z504567/1 and Biotechnology and Biology Research Council (BBSRC) projects BB/N01992X/1, BB/X008711/1 and BBSRC studentship BB/M011224/1 awarded to AC.

## Notes

### Competing Interest Statement

The authors have declared no competing interest.

### Summary of Updates

This version has been revised to include one missing author into the metadata.

