## Supplementary figures for "Enhancing cortical electrophysiological signatures of sleep overrides ER stress responses in sleep-deprived mice"

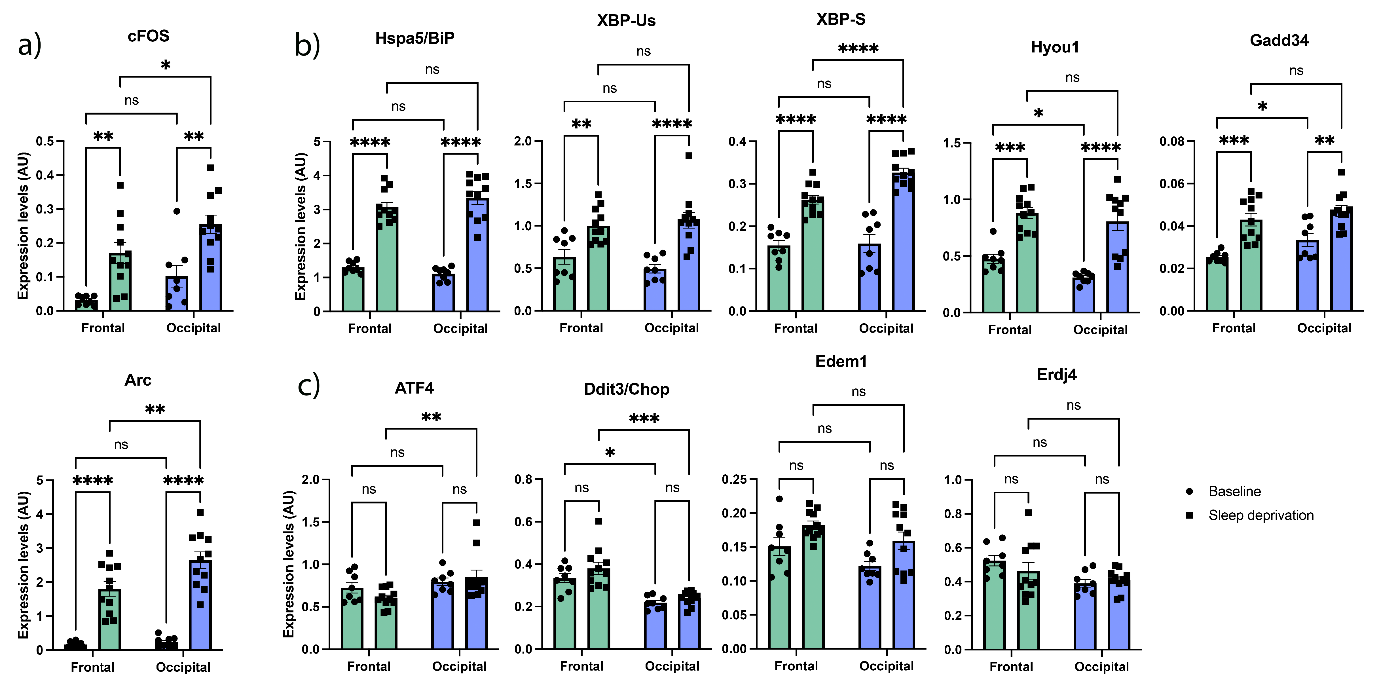

**Supplementary Figure 1: Activity-dependent and ER-stress related gene expression was also analysed by 2-way ANOVA for regional effects**. (A) The activity-dependent genes, *Fos* and *Arc*, showed a significant regional effect, with higher expression in occipital cortex after SD (2-way ANOVA region: *Fos* p=0.0021; *Arc* p=0.0070). (B) most of the ER-stress related genes that showed increased expression after SD displayed no significant regional variation with the exception of spliced *Xbp1* (2-way ANOVA region: Xbp1s p=0.0002). (C) Some ER-stress related genes that are not significantly upregulated by sleep deprivation, show variable regional expression.

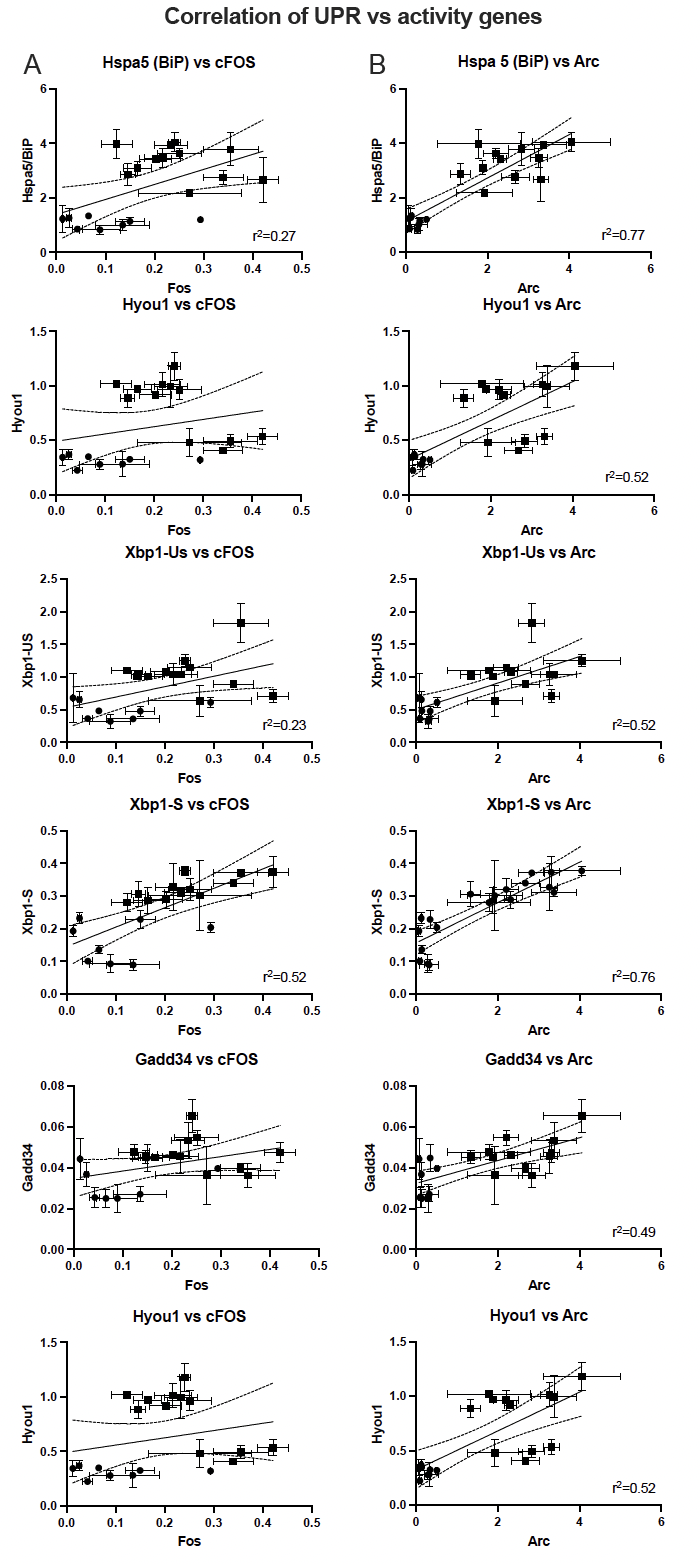

**Supplementary Figure 2: Sleep-deprivation causes positively correlated increases in ER-stress related and activity-dependent genes.** (A) Correlation between ER-stress related genes and *Fos* gene expression in occipital samples of cortex taken from control (baseline; circles) or 4h sleep-deprived (squares) mouse brains. *Hspa5* and *Xbp1* spliced and unspliced show significant positive correlation between ER-stress related gene expression levels and *Fos* expression levels. (B) Correlation between ER-stress related genes and *Arc* gene expression in occipital samples of cortex taken from control (baseline; circles) or 4h sleep-deprived (squares) mouse brains. All five genes show significant positive correlation between UPR gene expression levels and *Arc* expression levels.
